# Cost implications of a nationwide survey of schistosomiasis and other intestinal helminthiases in Sudan

**DOI:** 10.1101/865113

**Authors:** Mousab Siddig Elhag, Yan Jin, Mutamad Ahmad Amin, Hassan Ahmed Hassan Ahmed Ismail, Sung-Tae Hong, Haein Jang, Young-Ah Doh, Seungman Cha

**Affiliations:** Communicable and Non-Communicable Diseases Control Directorate, Federal Ministry of Health, Khartoum, Sudan; Department of Microbiology, Dongguk University College of Medicine, Gyeongju, South Korea; Research and Grants Unit, Ahfad University for Women, Omdurman, Khartoum, Sudan; Department of Parasitology and Tropical Medicine Seoul National University College of Medicine, Seoul, South Korea; Department of Development and Cooperation, Korea International Cooperation Agency, Sungnam, South Korea; Department of Global Development and Entrepreneurship, Graduate School of Global Development and Entrepreneurship, Handong Global University, Pohang, South Korea; Department of Disease Control, London School of Hygiene & Tropical Medicine, London, UK

**Keywords:** cost, a nationwide survey, Sudan, schistosomiasis, intestinal helminthiases

## Abstract

**Background:** It is vital to share details of concrete experiences of conducting a nationwide survey, so that the global health community could adapt it to expand geographic mapping programs, eventually contributing to the development of control and elimination strategies with limited resources. A nationwide survey of schistosomiasis and nine other intestinal helminthiases was conducted from December 2016 to March 2017 in Sudan.

**Objectives:** We aimed to describe details of the key activities and components required for the nationwide survey of schistosomiasis and other intestinal helminthiases and to analyze its costs.

**Methods:** We estimated financial and economic costs from the provider’s perspective. Cash expenditures incurred to implement the survey were defined as financial costs. We took into account all of the resources invested in the survey for economic costs, including the components that were not paid for, such as vehicles and survey equipment provided by the Ministry of Health, Sudan and the opportunity costs of primary school teachers’ time spent on the survey. We ran one-way sensitivity and probabilistic analyses using Monte-Carlo methods with 10,000 draws to examine the robustness of the primary analysis results.

**Results:** A total of USD 1,465,902 and USD 1,516,238 was incurred for the financial and economic costs, respectively. The key cost drivers of the nationwide survey were personnel and transportation, for both financial and economic costs. Personnel and transportation accounted for around 64% and 18% of financial costs, respectively.

**Conclusions:** The cost is expected to vary depending on the quantity and quality of existing laboratory facilities, equipment, and consumables, and the capability of laboratory technicians and sample collectors. Establishing central-level and independent supervision mechanisms to ensure the quality of the survey is equally important. We expect the global health community to draw on this study when developing nationwide surveys of schistosomiasis and other intestinal helminthiases.

**Author Summary:** Although large-scale mapping of schistosomiasis and other intestinal helminthiases has been conducted in some countries, little is known about the details of nationwide surveys, such as the necessary scale of the workforce, logistics, and the cost of conducting a nationwide survey. A nationwide survey of schistosomiasis and nine other intestinal helminthiases was conducted from December 2016 to March 2017 in Sudan. A total of 105,167 students participated in the survey from 1,772 primary schools in 183 districts of all 18 states of Sudan. Herein, we present the activities that were necessary to prepare and conduct a nationwide neglected tropical disease survey, along with details on the types and amounts of personnel, survey equipment, and consumables that are required. In addition, through an analysis of the costs of the nationwide survey, we generated average costs at the district and sub-district level. The key cost drivers were personnel and transportation, both of which were recurrent costs. Establishing a steering committee to develop and reach consensus on a survey protocol, assessing the capacities of potential staff (particularly laboratory technicians), and training laboratory technicians and data collectors were key components required to prepare a nationwide survey. If a government finds a way to mobilize existing government officials with no additional payment using the health system already in place, the cost of a nationwide survey would be remarkably lower. We expect the global health community to draw on this study to develop nationwide surveys for schistosomiasis and other intestinal helminthiases.

## Background

Neglected tropical diseases (NTDs) have been chronically underfunded, making it necessary to allocate limited resources efficiently [1, 2]. Although investments in disease mapping have increased in the past decade, in many parts of the world, the prevalence of NTDs remains unknown or patchy, or the data are outdated [3, 4]. The gaps in the known prevalence of schistosomiasis and intestinal helminthiases, including soil-transmitted helminthiases (STHs), are considerable, and prevalence data are only available for fewer than half of all districts in sub-Saharan Africa [5].

Implementation of NTD control has been hampered by a lack of data on their geographical distribution and limited funds. Operational experience with nationwide NTD surveys is limited, and this is particularly true for integrated surveys [6]. Therefore, it is vital to share the details of concrete experiences of conducting a nationwide survey, so that the global health community could adapt it to expand geographic mapping programs, eventually contributing to the development of control and elimination strategies with limited resources.

In recent years, several studies [6–10] have begun exploring the main drivers of NTD survey costs and the potential effects of modifying standard operating procedures. The first study [6] analyzed the cost of the integrated surveys for lymphatic filariasis, schistosomiasis, and STH infection conducted in South Sudan by Kolaczinski and colleagues [6]. They called on the global community to undertake similar studies to compare results across different settings. However, the survey in South Sudan was not conducted throughout the nation (instead, it was carried out only in one state), and previous studies [6–10] were limited to analyzing cost only. We need to understand what activities are necessary to prepare and conduct a nationwide NTD survey, and the types and amounts of personnel, survey equipment, and consumables that are required.

Sudan was one of the African countries with the largest unmapped areas of schistosomiasis and STHs, and it was necessary to update the prevalence data [11]. A nationwide survey of schistosomiasis and other intestinal helminthiases, including STHs, was conducted from December 2016 through March 2017, in which 105,167 students participated from 1772 schools in 183 districts of all 18 states of Sudan. Herein, we aim to describe the details of the key activities and components required to prepare and undertake a nationwide survey of schistosomiasis and other intestinal helminthiases, and to analyze the costs of the nationwide survey. We compare the results with those of previously conducted surveys to obtain insights into the key cost drivers of a nationwide survey of schistosomiasis and other intestinal helminthiases and to contribute to the development of survey designs elsewhere. Sharing experiences of nationwide surveys is important for the global health community to expand global mapping activities [12]. To the best of our knowledge, this is the first study of its kind to examine the details of the activities, logistics, and costs of a nationwide survey of schistosomiasis and other intestinal helminthiases.

## Methods

This nationwide survey was carried out under the umbrella of the SUKO project, which was the Sudan-Korea collaboration project of schistosomiasis and other intestinal helminthiases control supported by the Korea International Cooperation Agency (KOICA). The survey protocol has been described in detail elsewhere [13].

### Ethical statement

For this study, ethical approval was obtained from the Federal Ministry of Health, Sudan (FMOH/DGP/RD/TC/2016) and the Korea Association of Health Promotion (130750-20,164-HR-020). Prior to the survey, we provided the survey protocol to the Ministries of Health of all 18 states, including a description of the proposed activities. We explained the survey protocol to the head teachers, teachers, and schoolchildren. We obtained informed consent from the head teachers and schoolchildren in written form. We could not obtain informed consent from the parents due to the large sample size. Instead, schoolteachers informed the parents about the survey details through students and checked for parental consent before launching the survey. Data collection was undertaken using tablet PCs and the data were anonymized.

### Study area

Sudan is the third largest country in Africa, comprising 189 districts in 18 states. The estimated Sudanese population is 37.4 million in 2016, of whom 45.6% are children younger than 15 years and 3.9% are people aged above 59 years. The White Nile, the Blue Nile, and the River Nile flow through the country. Sixty-one percent of people have access to improved drinking water and 27% to improved sanitation. The study population was students aged 8-13 at 15,761 primary schools across the country.

### Sampling

We used two-stage random sampling for the nationwide survey. We applied probability-proportional-to-size sampling to select the schools. Twenty students from the second, fourth, and sixth grades were selected at each school. Based on compelling evidence of the focalized nature of schistosomiasis infections, we divided each district into one to three different ecological zones depending on its distance to water bodies (near, less than 1 km; medium, 1-5 km; far, 5 km or above). We defined an ecological zone as an area located within a similar distance from bodies of water in a district. There were only one or two ecological zones in some districts. We used random sampling for schools and students to derive precise estimates of prevalence with a sufficient sample size. Finally, we surveyed 105,167 students from 1772 primary schools from 390 ecological zones in 183 districts of 18 states across Sudan. The nationwide survey was conducted from December 2016 to March 2017.

### Detection of schistosomiasis and other helminthiases

Stool and urine samples were processed within 24 hours. A training module was given to each state level team, which included color images of the various parasites expected. We used the Kato-Katz technique for the eggs of *Schistosoma mansoni* and other intestinal helminthiases using two smears [14]. We used centrifugation to examine *S. haematobium* eggs. Two slides were observed for each sample and a federal supervisor cross-checked the results of the examinations by state laboratory technicians to validate them. Federal-level supervisors randomly selected 10% of slides examined by laboratory technicians and re-examined them on a daily basis. An independent team of biologists and parasitologists re-examined 5% of the slides for quality control purposes. A total of 655 people were deployed for the survey.

### Collection of data

For cost analysis, we applied the same methods suggested by Kolaczinski and colleagues [6]. We estimated financial and economic costs from the provider’s perspective. Cash expenditures incurred to implement the survey were defined as financial costs. We organized costs into capital and recurrent items. We calculated the average financial daily cost using straight-line depreciation for capital items. The total number of survey days was considered for the capital costs. We took into account all the resources invested in the survey, including the components that were not paid for, such as vehicles and survey equipment provided by the Ministry of Health, Sudan and the opportunity costs of primary school teachers’ time spent on the survey. Capital items were discounted with a 3% discount rate over their lifespan. Daily economic costs for all capital items were calculated and multiplied by the number of days they were used for the survey. We used the same estimated lifespan for vehicles and survey equipment (4 and 2 years, respectively) that were used in South Sudan [6], since similar harsh climactic conditions are present in both Sudan and South Sudan. Data for costs were collected from the financial records of the project. We used an average exchange rate of 1 USD = 0.15264 SDG for the costs of survey activities between September 2016 and March 2017. We did not use shadow prices. To estimate the total value of goods or services, we multiplied their unit price by the total number of each item. For office overhead costs, we directly requested the implementing organization to share the actual amount with the study team [15]. We presented the costs by district, sub-district, schools, and individual levels to allow for comparison with other studies. For key activities, real-time weekly and monthly reports were documented with budget components.

### Sensitivity analysis

We ran one-way sensitivity and probabilistic analyses using Monte-Carlo methods with 10,000 draws to examine the robustness of the primary analysis results [15]. Discount rates and the lifespan of vehicles and survey equipment were adjusted. A cumulative probabilistic curve was examined for the cost of survey per ecological zone and district.

## Results

Table 1 shows the main activities conducted in the nationwide survey from its preparation to results-sharing. A steering committee was established to supervise the overall program, consisting of high-profile Sudanese government officials, the World Health Organization country office staff, and parasitologists or epidemiologists from both Sudan and Korea. The first committee meeting was held in October 2016 to reach a consensus on the study protocol. After the committee agreed on the protocol, a workshop was organized to train state-level laboratory technicians and potential state-level supervisors. Thirty-six laboratory technicians and 18 state government officials were invited from 18 states. An additional objective of this workshop was to examine the capability of state-level laboratory technicians in order to help the management team understand the required intensity and duration of training to be conducted prior to the survey. During the three-day workshop, they were trained and their capability was assessed through classroom-based lectures and field-based mock surveys. Overall, the state-level laboratory technicians’ capability was found to be satisfactory. Most of them were laboratory technicians working for the state-run central laboratory, and they had experience examining various parasites for their routine work. Therefore, they were appropriately equipped with the required skills to detect schistosomiasis and other intestinal helminthiases through microscopy and centrifugation. All personnel for the survey were recruited in December 2016, including 18 federal-level supervisors for laboratory examinations, 18 federal-level supervisors for field visit activities for specimen collection and questionnaire administration, 18 state-level supervisors; 218 state-level laboratory technicians, 244 state level field-based data collectors, 3 independent quality control supervisors, 3 federal-level operation team members, and 2 independent data management support team members.

**Table 1.** Main activities and financial costs

| Time | Day | Main activities | Purpose | Trainers/Facilitators | Participants | Cost (US\$) | % |
| --- | --- | --- | --- | --- | --- | --- | --- |
| October 23~25, 2016 | 3 | Workshop | Examine the capacity of laboratory technicians and state coordinators | Senior laboratory technicians of Blue Nile Institute (1 Professor, 1 PhD)<br>Federal Ministry of Health (1 PhD, 1 MD)<br>External specialists (2 Professors, 1 PhD)<br>Project management staff (5) | State level laboratory technicians (36)<br>State coordinators (18) | 17,750 | 1.1% |
| October 23, 2016 | 1 | Steering board meeting | Reach a consensus on the study protocol of a nationwide survey | Project management staff (5) | Board members (8) | 1,448 | 0.1% |
| December 18~22, 2016 | 5 | Workshop & Training | Orientation<br>Introduce overall project<br>Help to understand study methods<br>Train data collection methods<br>Train laboratory examination | Federal Ministry of Health (1 PhD, 1 MD)<br>External specialists (2 Professors, 1 PhD, 1 MSc)<br>Project management staff (5) | State supervisors (18)<br>Federal Supervisors - Senior laboratory technicians (18)<br>- Supervisors for specimen collection and household survey (18) | 64,791 | 3.9% |
| December 31, 2016- January 10, 2017 | 1 | Workshop | Introduce project; Obtain consent | State Coordinators | Head teacher from target schools (1811) | 92,612 | 5.6% |
| December 31, 2016 ~ March 15, 2017 | 12-30 | Nationwide survey | Carry out a nationwide survey on schistosomiasis and other helminthiases | State level laboratory technicians (218)<br>State level data collectors (244)<br>Federal level supervisors (36)<br>Independent quality control supervisors (3)<br>External specialists (1)<br>Central level operation team (5)<br>Independent data management support team (2) | 105,167 students, 1772 schools in 390 ecological zones, 183 districts | 1,465,902 | 88.4% |
| March 19<br>2017 | 1 | Post-survey<br>workshop | Share lessons learnt<br>Collect suggestions<br>and<br>recommendations for<br>a next round of<br>nationwide survey<br>Discuss details of a<br>community-based<br>survey | Federal Ministry of Health (1 PhD, 1 MD)<br>External specialists (2 Professors, 1 PhD)<br>Project management staff (5) | State coordinators (18)<br>Federal Supervisors<br>- Senior laboratory<br>technicians (18)<br>- Supervisors for specimen<br>collection and household<br>survey (18) | 10,745 | 0.6% |
| March 20<br>2017 | 1 | Steering board<br>meeting | Share key findings<br>Understand policy<br>implications<br>Discuss future plans | Project management staff (5) | Board members (8) | 5104 | 0.3% |
| Grand<br>Total |  |  |  |  |  | 1,658,352 | 100% |

In December 2016, 36 federal-level supervisors and 18 state-level supervisors were trained for five days in Khartoum, the capital city. It was ensured that they understood the survey objectives, student sampling methods, and details of questionnaires. They practiced laboratory techniques at National Laboratory of Federal Ministry of Health, Sudan to examine samples for *S. haematobium, S. mansoni*, and nine other intestinal helminthiases, including STHs, using real parasite egg samples. A private consultant company was contracted to develop and install a program to facilitate the real-time management of data. The consultant trained the supervisors on how to enter and manage data with tablet PCs purchased by the project team. Prior to the survey, the head teachers of 1811 target primary school were invited by the Ministry of Health at the state level. State-level supervisors helped them to understand the overall objectives and methods of the study, and obtained their consent with, and support for, the survey between late December 2016 and early January 2017. The state-level laboratory technicians and data collectors were trained by federal- and state-level supervisors. Thirty-six federal-level supervisors were deployed to each state for the entire period of the nationwide survey, during which they undertook daily monitoring and supervision, and reported real-time results to the central operation team.

The nationwide survey was conducted between December 31, 2016 and March 15, 2017. The starting and ending date varied depending on the state, and the duration of the survey in each location ranged from 12 days to 30 days. Immediately after the survey, in March 2017, federal- and state-level supervisors participated in a post-survey workshop conducted to collect and share lessons learned and experiences, and to make recommendations for future surveys. Finally, a steering committee meeting was held to share key findings, discuss future plans, and derive policy recommendations. A total of USD 1,658,352 (financial cost) was spent for the overall activities of the nationwide survey, including preparation, the post-survey workshop, and steering committee meeting, of which USD 1,465,902 (88%) was incurred by the actual survey.

A reference case scenario is outlined in S1 Appendix, in which we applied the same scenario that was used for costing in South Sudan [6]. Table 2 shows the total financial and economic costs for the nationwide survey. A total of USD 1,465,902 and USD 1,516,238 was incurred for the financial and economic costs, respectively. The key cost drivers of the nationwide survey were personnel and transportation, for both financial and economic costs. Personnel and transportation accounted for around 64% and 18% of financial costs, respectively. Capital costs amounted to only 0.5% of the total financial costs and 1.4% of the total economic costs. There was a slight increase in capital economic costs, which mainly resulted from the opportunity cost of vehicles and survey equipment supported by the Ministry of Health, not paid for by the SUKO project. The fuel for those government vehicles was covered by the SUKO project and was already included in the financial costs. Seventeen government vehicles and 144 microscopes were provided by the Ministry of Health, Sudan with no charge for the SUKO project. Tables 3 and 4 present the details of financial and economic costs, respectively. For the survey, 422 vehicles × days and 3,464 microscopes × days were supported, which resulted in USD 10,886 and USD 2,877, respectively. In total, 5,316 school teachers were mobilized to facilitate specimen collections and questionnaire-based surveys; at one day per teacher, this was translated into USD 94,811 of economic costs. By implementation unit, the economic cost was USD 8,285 per district or USD 3,888 per ecological zone. Based on the results of sensitivity analyses adjusting for the discount rate or lifespan of vehicles, the estimates did not change to a meaningful extent (Table 5, Figure 1). The results of the Monte Carlo analysis with 10,000 trials show a range of all the cost metrics. Figure 1 presents the cumulative density functions of the costs at the district and sub-district levels. The fifth percentile of the costs was US$ 8,215 and USD 3,855 at the district and sub-district level, respectively, while the 95th percentile reached USD 8,357 and USD 3,921.

**Fig 1.**
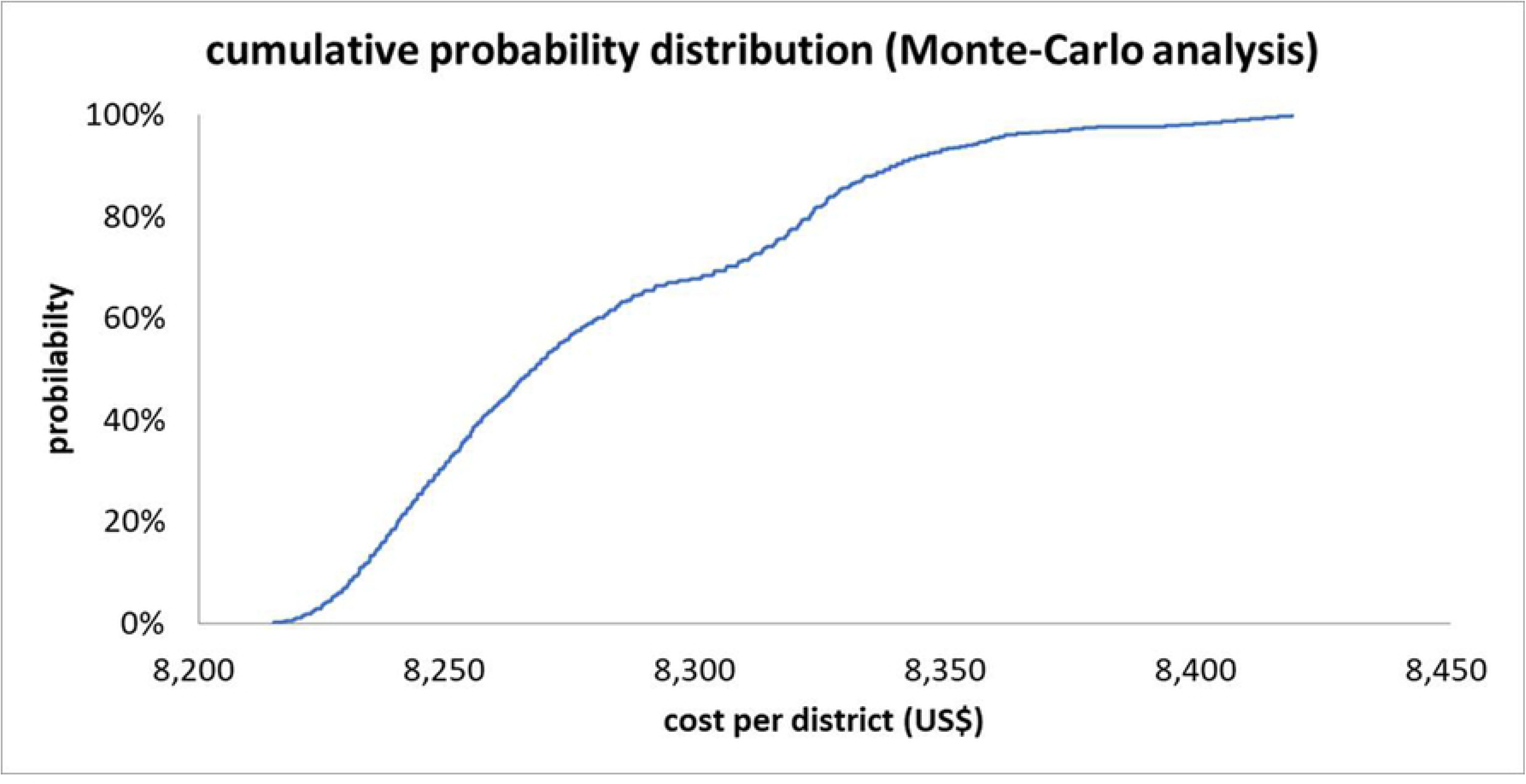
Cumulative probability distribution (Monte-Carlo probabilistic sensitivity analysis, 10,000 draws)

**Table 2.** Financial and economic costs

|  | Total financial cost |  | Total economic cost |  |
| --- | --- | --- | --- | --- |
| | US\$ | % | US\$ | % |
| **Capital cost** |  |  |  |  |
| survey equipment | 7,256 | 0.5% | 10,461 | 0.7% |
| Vehicles |  |  | 10,886 | 0.7% |
| Sum | 7,256 | 0.5% | 21,347 | 1.4% |
| **Recurrent cost** |  |  |  |  |
| Transportation (vehicle hiring) | 260,479 | 17.8% | 260,479 | 17.0% |
| Personnel | 942,272 | 64.3% | 978,517 | 64.0% |
| Survey consumable | 54,687 | 3.7% | 54,687 | 3.6% |
| Others | 170,104 | 11.6% | 170,104 | 11.1% |
| Sum (recurrent cost) | 1,427,542 | 97.4% | 1,463,787 | 95.7% |
| Total (capital and recurrent cost) | 1,434,798 | 97.9% | 1,485,134 | 97.1% |
| Overhead | 31,104 | 2.1% | 31,104 | 2.0% |
| Grand | 1,465,902 | 100.0% | 1,516,238 | 99.1% |
| District | 8,010 |  | 8,285 |  |
| Ecological zone | 3,759 |  | 3,888 |  |
| School | 827 |  | 856 |  |
| Person | 14 |  | 14 |  |

**Table 3.**
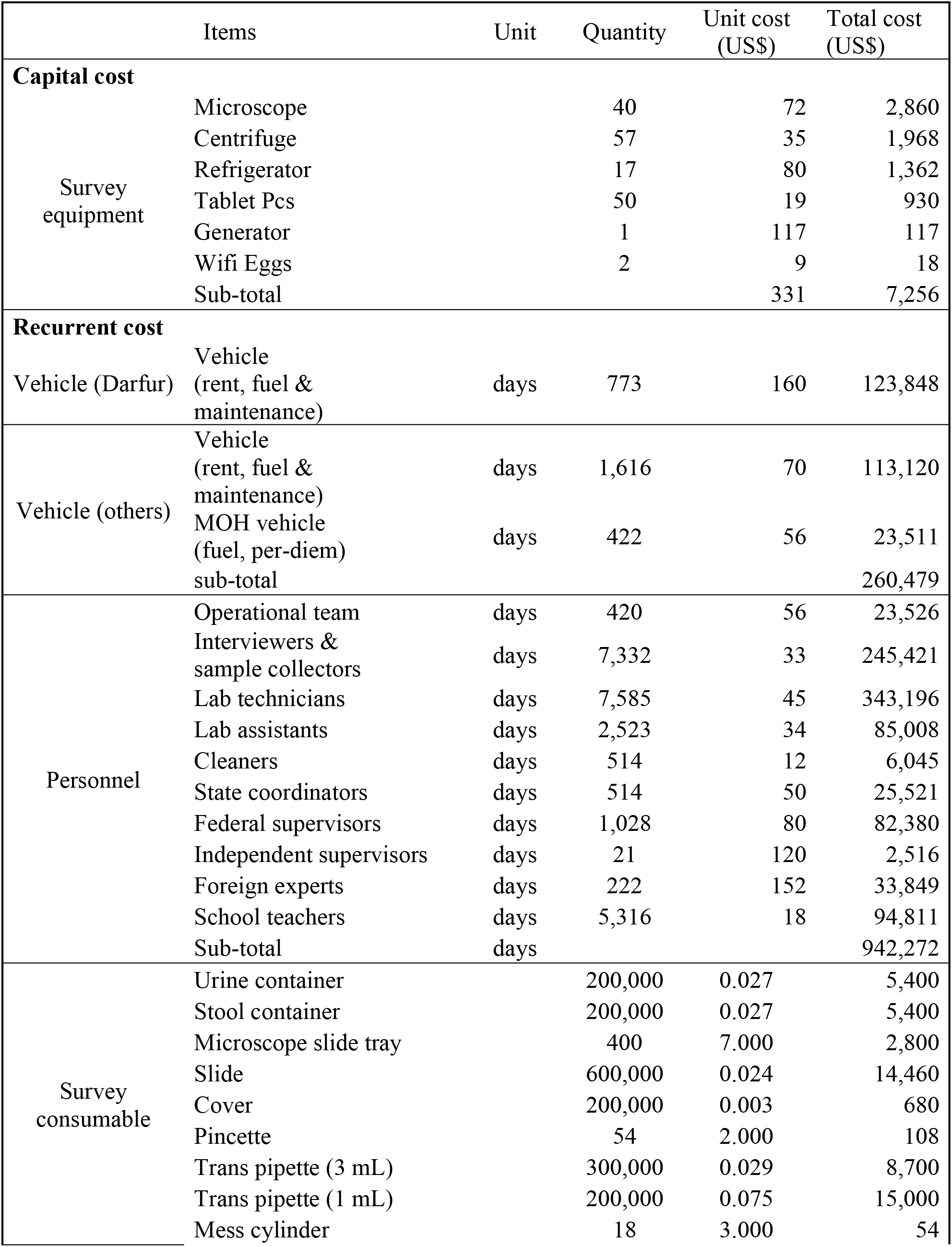
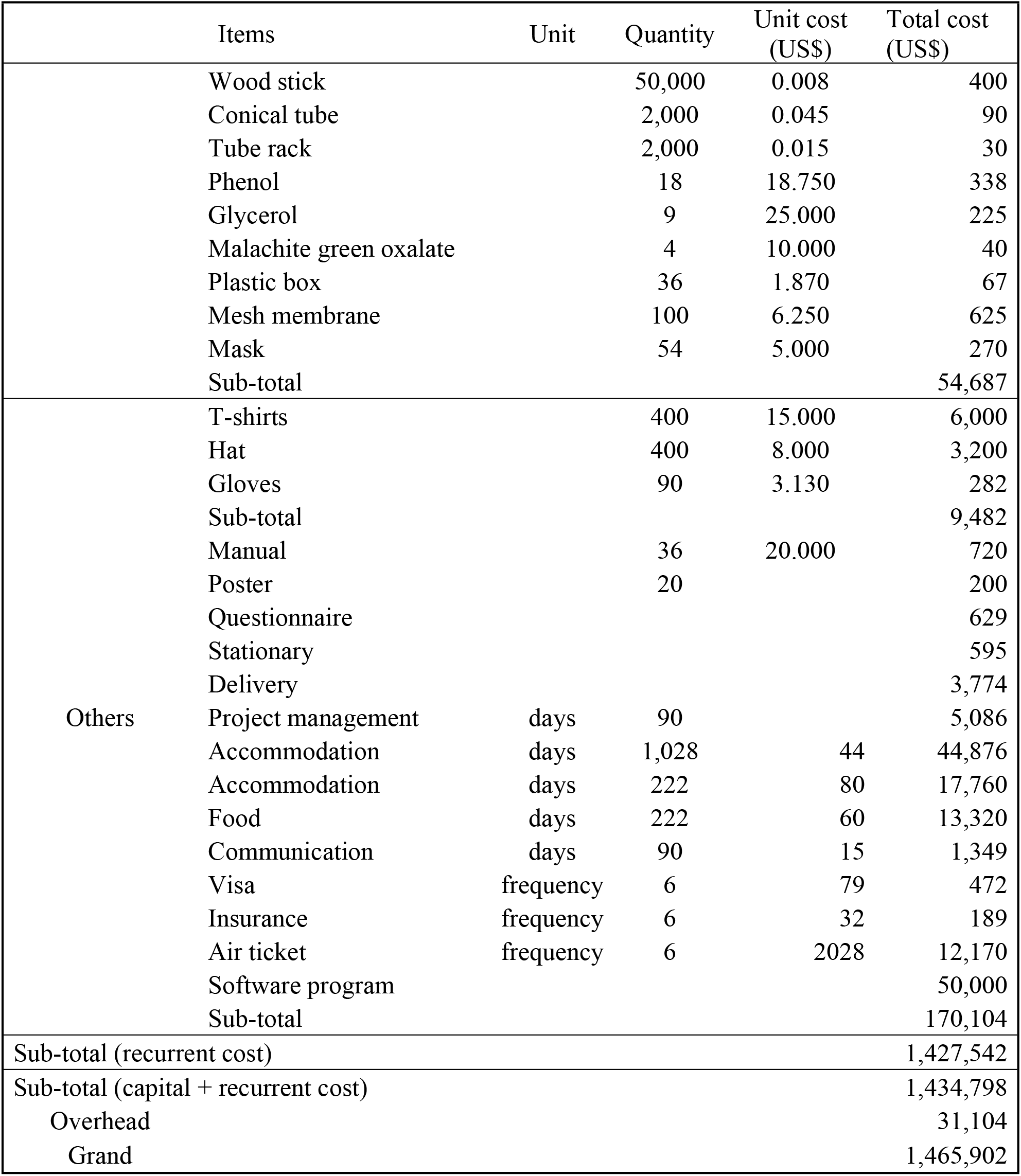
Details of financial costs

**Table 4.**
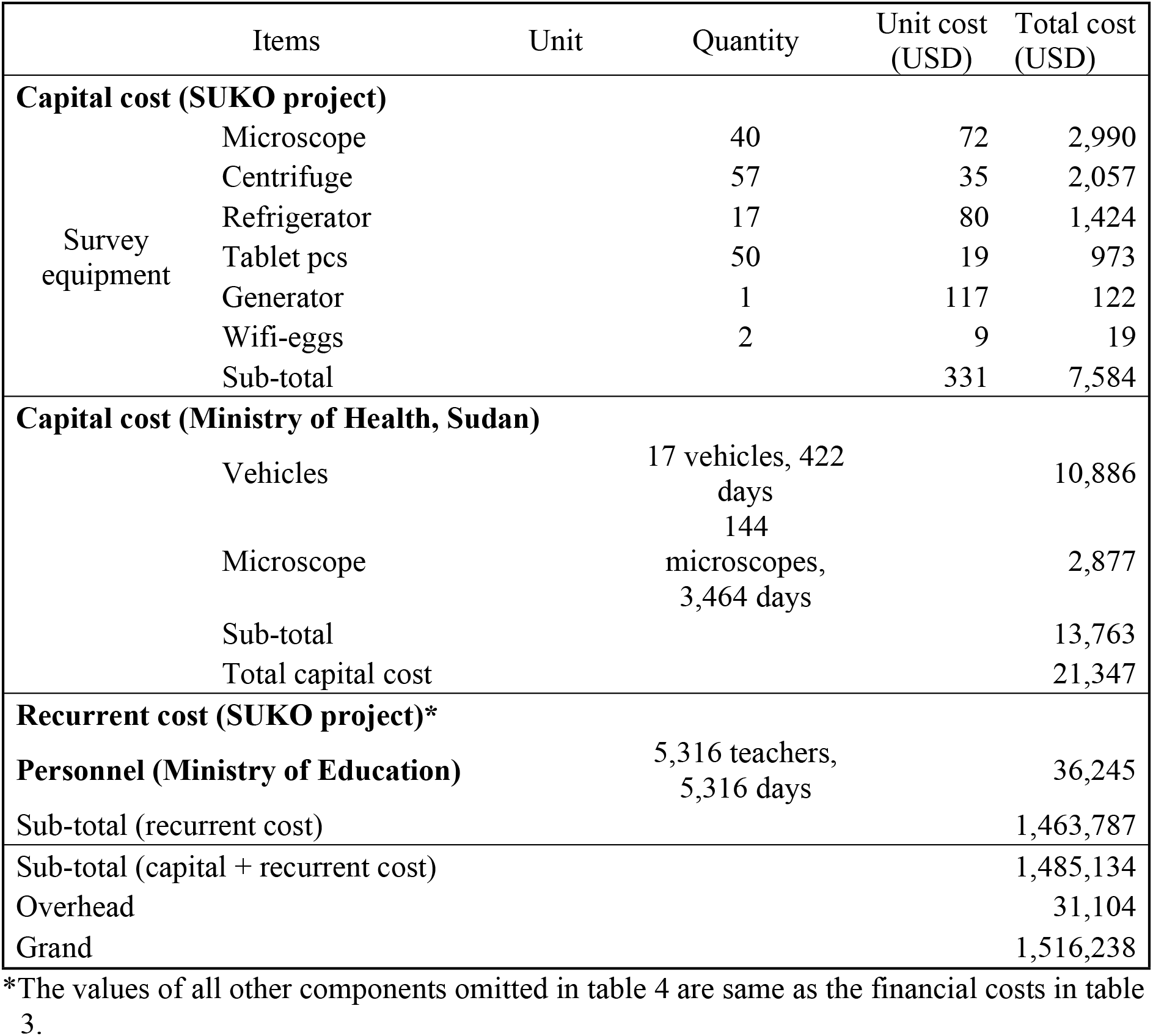
Details of economic costs

**Table 5.** Sensitivity analysis to cost estimates

| Assumptions tested |  |  |  | District level | % Deviation from base case | Ecological zone level | % Deviation from base case |
| --- | --- | --- | --- | --- | --- | --- | --- |
| **One-way** | Base case |  |  | 8,285 |  | 3,888 |  |
|  | Discount rate | 1% |  | 8,282 | -0.04% | 3,886 | -0.05% |
|  |  | 10% |  | 8,303 | 0.22% | 3,896 | 0.21% |
|  | Vehicle lifespan | 8 years |  | 8,258 | -0.33% | 3,875 | -0.33% |
|  | Equipment lifespan | 4 years |  | 8,259 | -0.31% | 3,875 | -0.33% |

## Discussion

We provided details of the nationwide survey of schistosomiasis and other intestinal helminthiases in terms of the key activities and costs in Sudan, therefore when planning to undertake future nationwide NTD surveys, the global health community could adapt and refine the activities and cost elements of the survey described herein.

The main cost drivers were recurrent costs: transportation (particularly hiring vehicles) and personnel. Prior to the survey, we assessed the capabilities of state-level laboratory technicians and data collectors. We invited 36 laboratory examiners and 18 state coordinators, who were purposively selected by the State Ministry of Health. Based on the assessment, we realized that there was a sufficient number of laboratory technicians qualified for the survey. If there had not been enough laboratory technicians with the required skills to detect schistosomiasis and other intestinal helminthiases through microscopy and centrifugation, the project team would have organized a more intensive training for a longer duration with closer supervision, all of which would have increased costs.

Additionally, Sudan had state-level laboratory facilities, many of which had microscopes and centrifuges, although the quantities were not sufficient. In some exceptional cases, such as North Kordofan, we had to operate a mobile laboratory due to the remoteness of some schools, but in general, laboratory work was carried out at state-run central laboratories. If a nationwide survey is to be conducted in settings with no local laboratory, more costs will be incurred for the laboratory setting, including the cost for equipment.

When designing the study protocol, the project team planned to purchase seven vehicles (six for the state-level survey team and one for the central-level operation team). However, vehicle purchase was not possible during the survey period, and therefore vehicles had to be hired from local rental-car companies, which led to a substantial rise in recurrent costs.

The reference case suggested by Kolaczinski and colleagues [6] was applied for this study. The nationwide survey of schistosomiasis and other intestinal helminthiases in Sudan was much less expensive than the survey conducted in 2009 in South Sudan, its neighboring country. The average economic cost of the schistosomiasis, STH, and lymphatic filariasis survey in South Sudan was USD 40,206 or USD 9,573 per district (county) or sub-district (*payam*), respectively. In Sudan the average economic cost was USD 8,285 or USD 3,888 respectively. In Sudan, the average economic cost per school or per student was USD 856 or USD 14, respectively. The results of the Monte-Carlo analysis demonstrated that the costs varied little across a range of discount rates and lifespans of vehicles and survey equipment (5–95th percentile of cumulative distribution: US$ 8,215-8,357 at the district level and US$ 3,855-3,921 at the sub-district level).

Caution is needed when interpreting the results of this study because our findings cannot be generalized to every other context with a high prevalence of schistosomiasis and other intestinal helminthiases, which is the major limitation of this study. In particular, Sudan has a number of highly skilled laboratory technicians, and NTDs are prioritized by both the Federal and State Ministry of Health [16, 17], making it feasible to mobilize a considerable number of capable workforce for conducting the survey. Another limitation of this study lies in the number of samples, which also relates to the generalizability of the study. Unlike the conventional methods recommended by the WHO making a decision of mass drug administration at the district level [18], we divided each district into smaller areas according to the proximity to water bodies to reflect the focalized nature of schistosomiasis prevalence [19–21]. In addition, we sampled a sufficient number of students to derive precise estimates of prevalence at the state level [22]. This led to sampling far more students at the district level than would have been done using the conventional method (250 students from five schools per district using conventional methods vs. 900 students from 15 schools per district in this study). If this survey had applied the conventional methods, the average economic cost per district would have amounted to the current cost per ecological zone of this study (i.e., USD 3,888).

Personnel (64%) and transportation (18%) were the key cost drivers in Sudan, while personnel (38%) and survey consumables (27%) were the key drivers in South Sudan [6]. Whereas the survey in South Sudan was conducted only in a single state [23], the survey in Sudan was undertaken in all 18 states, which required an enormous workforce and a large amount of vehicles. Additionally, for the nationwide survey in Sudan, we incorporated federal-level supervision and an independent supervision mechanism for enhancing quality control, which led to a rise in costs. The participants of this study were 105,167 students from 1,772 schools in 390 ecological zones in 183 districts of all 18 states of Sudan. We suggest that the greatest share of costs for personnel and transportation should be reflected when budgeting future surveys for schistosomiasis and other intestinal helminthiases.

Apart from the key activities and costs, we provided considerable detail about the nationwide survey including the duration of the survey, the number of personnel by position and role, and the total number of each capital or recurrent item. We estimated the costs from the provider’s perspective. However, if we had applied the societal perspective for costing [24, 25], the result would not have changed considerably because few items were not paid for in this survey. No volunteers [26] were mobilized and parents or community members were not participated in this survey because it was conducted at the school level. We used the same lifespan for capital items as in the reference case in the South Sudan [6]. The relatively short lifespan of capital items (i.e., 4 years for vehicles and 2 years for survey equipment) made the cost estimates more conservative.

## Conclusion

Establishing a steering committee to develop and consensus a survey protocol, assessing the capabilities of potential staff (particularly laboratory technicians), and training laboratory technicians and data collectors are key components required to prepare a nationwide survey. Collecting lessons learned from frontline staff, sharing the results, and developing future plans with stakeholders are crucial activities after carrying out a nationwide survey. Key cost drivers are personnel and transportation. In particular, the personnel component accounts for a substantial share of the cost. The cost is expected to vary depending on the quantity and quality of existing laboratory facilities, equipment, and consumables, and the capability of laboratory technicians and sample collectors. Establishing central-level and independent supervision mechanisms to ensure the quality of the survey is equally important. It is worth noting that if a government finds a way to mobilize existing government officials with no additional payment using the health system already in place, the cost of a nationwide survey would be remarkably lower. To the best of our knowledge, this is the first empirical study to investigate the key activities and cost drivers of a nationwide survey of schistosomiasis and other intestinal helminthiases. We expect the global health community to draw on this study when developing nationwide surveys of schistosomiasis and other intestinal helminthiases.

## Acknowledgement

This project was supported by the Korea International Cooperation Agency (KOICA) under the “Integrated schistosomiasis and soil-transmitted helminthiasis control program, Sudan (00145)”. The authors thank the project team members for their efforts and contributions to controlling neglected tropical diseases in Sudan. They extend their appreciation to community members, the Ministries of Heath of 18 states, and the Federal Ministry of Health, Sudan. Special thanks go to Dr Nahid Abdelgadir, International Health Directorate Project Management Unit, the Federal Ministry of Health, Sudan.

## Supporting information

S1. Appendix. Reference case scenario

